# Efficient indexing of peptides for database search using Tide

**DOI:** 10.1101/2022.09.30.510396

**Authors:** Frank Lawrence Nii Adoquaye Acquaye, Attila Kertesz-Farkas, William Stafford Noble

## Abstract

The first step in the analysis of protein tandem mass spectrometry data typically involves searching the observed spectra against a protein database. During database search, the search engine must digest the proteins in the database into peptides, subject to digestion rules that are under user control. The choice of these digestion parameters, as well as selection of post-translational modifications (PTMs), can dramatically affect the size of the search space and hence the statistical power of the search. The Tide search engine separates the creation of the peptide index from the database search step, thereby saving time by allowing a peptide index to be reused in multiple searches. Here we describe an improved implementation of the indexing component of Tide that consumes around four times less resources (CPU and RAM) than the previous version and can generate arbitrarily large peptide databases, limited by only the amount of available disk space. We use this improved implementation to explore the relationship between database size and the parameters controlling digestion and PTMs, as well as database size and statistical power. Our results can help guide practitioners in proper selection of these important parameters.

## 1 Introduction

Every mass spectrometry protein database search engine must solve two primary tasks: producing from a given protein database a corresponding list of peptides, and scoring each observed fragmentation spectrum relative to peptides in the list. In some search engines, these two functions are tightly coupled. In Comet, for example, each protein is extracted sequentially from the database, digested to peptides, and then scored against an observed spectrum [1]. In contrast, Tide separates the search into two phases: peptide indexing and searching [2]. The peptide indexing step happens before any of the spectra are scored, and the index of peptides is stored on disk. This approach saves computation when the same database will be searched multiple times. Also, by storing unique peptides in the index, the scoring phase avoids re-scoring any peptide that recurs in multiple proteins.

The initial description of the Tide search engine focused on optimizing the search component [2]. As a result, until now the indexing step of Tide (implemented in the Crux toolkit [3] as the command `tide-index`) has been relatively slow. Furthermore, because the naive implementation involves sorting all of the peptides in memory, any user interested in a very large database—e.g., by considering large numbers of post-translational modification (PTMs), relaxed enzymatic digestion rules, or searching large metaproteomic databases—needed to have very large amounts of memory available for the indexing step.

Here we describe an optimized version of tide-index that dramatically reduces the time and memory requirements of the tool. First, we restructured the core data structures and reorganized the code so that the components work together efficiently. This was required because the tide-index code had become fragmented during implementation of new features by different developers. As a result of these optimizations, tide-index is now 3–5 times faster and consumes around 4 times less memory than its predecessor. Second, we modified tide-index so that sorting and storing the unique peptides can be done using external disk operations, subject to a user-specified parameter called `memory-limit`. In this way, tide-index can now generate very large peptide database files. For instance, we used tide-index to generate an index containing 67.4 billion unique peptides from the human proteome, using non-enzymatic digestion and allowing three commonly used variable modifications per peptides. This operation required nearly five days, and the resulting index occupies 4 TB of disk space. In principle, tide-index is able to generate arbitrarily large peptide index files, limited only by the capacity of the disk storage.

### Box 1: Parameters controlling enzymatic digestion and PTMs.

- The **digestion rule** specifies where the protease cleaves. For example, trypsin canonically cleaves after K or R residues, and chymotrypsin cleaves after F, W, Y, or L. In this work, we focus solely on tryptic digestion, though tide-index allows for any type of digestion rule.
- The **digestion** parameter controls whether cleavage occurs only at the sites specified by the digestion rule (full digestion), occasionally also at sites that are not specified by the rule (partial digestion), or anywhere in the sequence (non-enzymatic digestion). This parameter can have a dramatic effect on the number of peptides produced by a single protein (Figure 3A).
- The **missed cleavages** parameter puts a limit on the number of digestion sites that can be included within a single peptide in the index (Figure 3B).
- A **static modification** is a PTM that is applied to every instance of a given amino acid in the database. The canonical static modification is carbamidomethylation of cysteine, which induces a mass shift of 57.021 Da. Because this type of modification occurs on every cysteine, the parameter has no effect on database size.
- A **variable modification** is a PTM that may or may not occur on each peptide. For example, phosphorylation induces a shift of 79.966 Da on serine, threonine, and tyrosine residues. A peptide that contains an amino acid subject to variable modification must be included twice in the peptide index, with and without the PTM applied. Consequently, a peptide with *n* modifiable amino acids gives rise to 2^*n*^ modified forms. For example, oxidation of methionine yields four variants of the peptide on the peptide MPEPMPEPK: MPEPMPEPK, M*PEPMPEPK, MPEPM*PEPK and M*PEPM*PEPEK (where * indicates oxidation).
- The **maximum number of modifications** (max-mods) parameter specifies the number of distinct amino acids that can harbor a PTM within a single peptide sequence.

We then used the optimized version of tide-index to systematically explore the relationship between various search parameters, database size, and statistical power. In particular, we aimed to provide practical insight into the impacts of parameters that control enzymatic digestion and PTMs (summarized in Box 1 and Figure 1). Using data from four different species, we quantify the impact of three parameters—the type of digestion, the number of missed cleavages, and the number of PTMs—on database size and the number of detected peptides, with and without inclusion of the Percolator machine learning post-processor [4]. The new version of Tide is now available in the Crux toolkit [3], which is available as Apache licensed open source code and as pre-compiled binaries for Linux, MaxOS and Windows at http://crux.ms.

**Figure 1:**
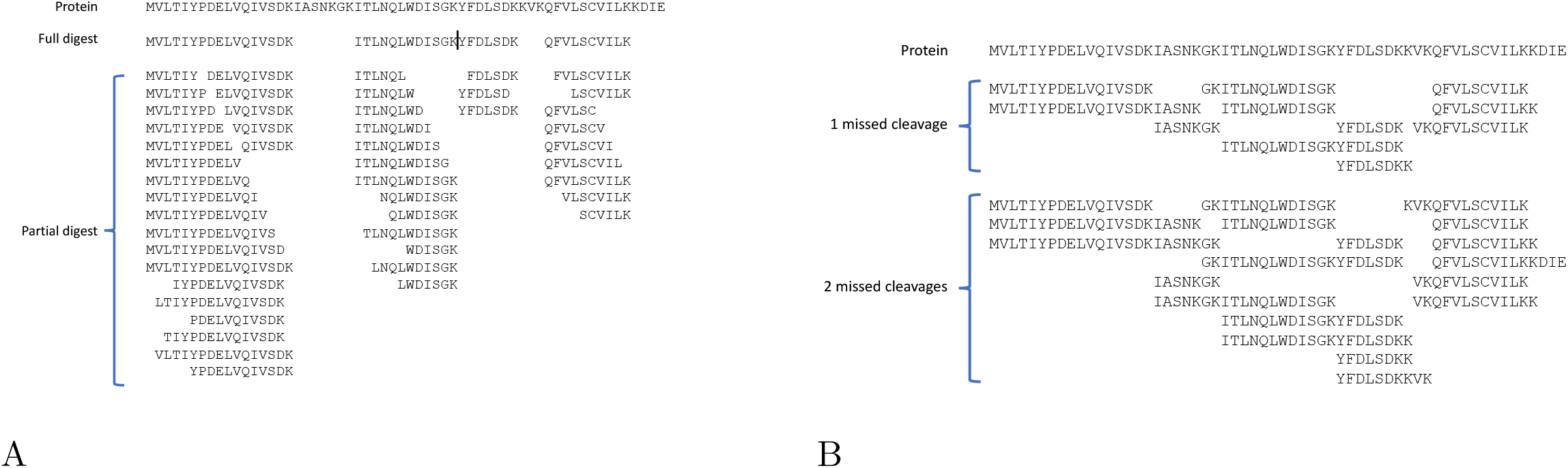
Examples of digestion rules. (A) Effect of digesting a simple protein sequence with trypsin, using full digestion or partial digestion. Peptides are limited to 7–50 amino acids in length. Non-enzymatic digestion (not shown) yields 1485 peptides for this protein. (B) Effect of digesting the same protein sequence with fully tryptic digestion but allowing one or two missed cleavages.

## 2 Methods

### 2.1 Optimizations for memory usage and performance

#### Memory

We began by modifying tide-index so that the user could control how much memory the program requires. Conceptually, tide-index’s operation is relatively straightforward. The program digests the given proteins to peptides, sorts them and discards redundant peptides, and then augments the resulting list of unique peptides with modified forms based on the user-specified list of PTMs. The original tide-index implementation stored and sorted all of the peptides in memory simultaneously. This step required a large amount of RAM for even relatively small sets of peptides. We therefore re-implemented the sorting-and-filtering routine to optionally make use of external disk operations. The new `memory-limit` parameter allows the user to specify the maximum amount of memory to be used for peptide filtering. Once the peptide container exceeds the memory limit during the peptide generation step, tide-index sorts and then dumps the peptides from the container to disk in a temporary file. After all peptides are generated, the temporary files are merged to create a final peptide index. We note that the `memory-limit` parameter introduces a trade-off between speed and memory usage: a low value of `memory-limit` leads to many small temporary files, which in turn requires more time for the merging step.

#### Speed

Since Tide’s initial publication in 2011, the tide-index code has been extended with various new features, which were implemented by different developers. In the process, the tide-index code became fragmented, the code complexity increased, and the code maintenance became cumbersome. At the same time, the code modifications led to an overall decrease in Tide’s efficiency. To address this problem, we restructured the tide-index code and optimized the underlying data structures so that all components in tide-index work smoothly together. For instance, we removed a container variable that unnecessarily stored an extra copy of each peptide. We also introduced new variables to link target and decoy peptides directly, thereby removing a subsequent step in which targets and decoys were paired together. This change eliminated an *O*(*n* log *n*) sort operation for *n* peptides.

### 2.2 Data

To test the new tide-index and to investigate the relationship between database size and statistical power, we downloaded one randomly selected raw spectrum files from each of five PRIDE data sets, representing a variety of organisms and instrument types (Table 1). For each data set, the precursor window size and PTM settings were taken from the associated papers. Carbamidomethylation of cysteine was specified as a fixed modification for all searches.

**Table 1:** Data sets. The table lists the five PRIDE datasets used in this study. “Precursor” indicates the precursor window size (in ppm), and “Fragment” is the fragment bin width (in Da). PTMs correspond to oxidation of methionine (O(M)), phosphorylation of serine, threonine and tyrosine (P(STY)), N-terminal acetylation (A(nT))), and TMT-6plex tags. For each PRIDE ID, one raw file was selected at random, listed on the second row of each table entry.

| PRIDE ID<br>File name | Species | Instrument | Precursor | Fragment | PTMs | Spectra |
| --- | --- | --- | --- | --- | --- | --- |
| PXD001928 [5]<br>Yeast_XP_Tricine_trypsin_147.mgf | *S. cerevisiae* | LTQ Orbitrap Velos | 20 | 1.0005079 | O(M) | 26,060 |
| PXD020052 [6]<br>PhDE170503_3D7icam_SCX3_F20584.mzML | *P. falciparum* | Orbitrap Fusion | 5 | 1.0005079 | O(M), A(nT) | 68,790 |
| PXD028806 [7]<br>20180913_Cell_Line_TMT_Test_F24_1.mzML | *H. sapiens* | Q-Exactive Plus | 10 | 0.02 | O(M), TMT | 61,594 |
| PXD028808 [8]<br>210312_A_MM_Aachen_14_TI0018_DIS_02_GH5_1_1958.mzML | *A. thaliana* | Impact II Q-TOF (Bruker) | 10 | 0.02 | O(M), A(nT) | 51,771 |
| PXD017407 [9]<br>20190304_231_all.mzML | *H. sapiens* | Q Exactive HF-X | 10 | 0.02 | O(M), P(STY) | 44,119 |

Reference proteomes of four different species, with and without isoforms, were downloaded from Uniprot on March 22, 2022. For reference, we include statistics about both versions of each reference proteome (Table 2), but all experiments in this paper were carried out using the canonical versions.

**Table 2:**
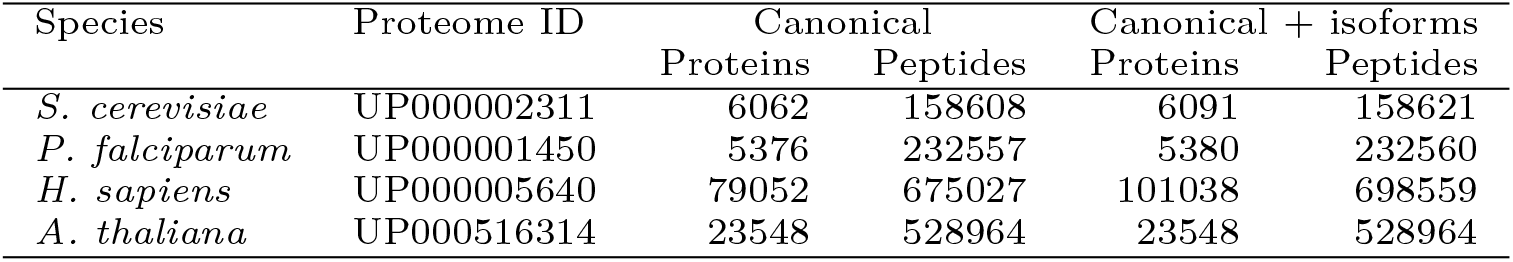
Reference proteome databases. The “peptides” column lists the number of peptides using full tryptic digestion with no missed cleavages allowed and without variable modifications. For reference, we include statistics about both versions (with and without isoforms) of each reference proteome, but all experiments in this paper were carried out using the canonical versions.

| Species | Proteome ID | Canonical |  | Canonical + isoforms |  |
| --- | --- | --- | --- | --- | --- |
|  |  | Proteins | Peptides | Proteins | Peptides |
| *S. cerevisiae* | UP000002311 | 6062 | 158608 | 6091 | 158621 |
| *P. falciparum* | UP000001450 | 5376 | 232557 | 5380 | 232560 |
| *H. sapiens* | UP000005640 | 79052 | 675027 | 101038 | 698559 |
| *A. thaliana* | UP000516314 | 23548 | 528964 | 23548 | 528964 |

To examine the impacts of peptide generation parameters and thus database size on the statistical power, we created a series of parameter settings (Table 3). The parameters vary enzymatic digestion, number of PTMs per peptide, and the maximum number of missed cleavages. The resulting series of peptide datasets is roughly increasing in the corresponding numbers of peptides.

**Table 3:** Parameter settings. All peptide data sets included decoy peptides generated by peptide reversal. The “Missed” column indicates the maximum number of missed cleavages, and “PTMs” indicates the maximum number of PTMs per peptide. The “Peptides” column indicates the number of peptides generated from the human proteome, and the “Time” column shows the execution time the new tide-index took. Note that Parameter setting #14 was only used for the immunopeptidomics experiment (Section 3.4), and additionally the minimum and maximum peptide lengths were set to 8 and 15, and TMT 6-plex labeling was added to the peptide n-terminal and lysine resides as static modifications.

| ID | Digestion | Missed | PTMs | Mods-spec | Peptides | Time (s) |
| --- | --- | --- | --- | --- | --- | --- |
| 1 | full-digest | 0 | 0 | - | 1,349,379 | 4 |
| 2 | full-digest | 0 | 1 | 1M+15.9949 | 1,784,632 | 6 |
| 3 | full-digest | 1 | 1 | 1M+15.9949 | 4,797,281 | 16 |
| 4 | full-digest | 2 | 1 | 1M+15.9949 | 8,169,434 | 26 |
| 5 | full-digest | 3 | 1 | 1M+15.9949 | 11,396,220 | 38 |
| 6 | full-digest | 4 | 1 | 1M+15.9949 | 14,198,510 | 46 |
| 7 | full-digest | 0 | 1 | 3M+15.9949, 3STY+79.966331 | 5,174,005 | 17 |
| 8 | full-digest | 0 | 2 | 3M+15.9949, 3STY+79.966331 | 13,030,652 | 54 |
| 9 | full-digest | 0 | 3 | 3M+15.9949, 3STY+79.966331 | 28,304,095 | 154 |
| 10 | full-digest | 3 | 3 | 3M+15.9949, 3STY+79.966331 | 300,423,422 | 3,380 |
| 11 | partial-digest | 0 | 1 | 1M+15.9949 | 34,021,904 | 103 |
| 12 | partial-digest | 3 | 3 | 3M+15.9949, 3STY+79.966331 | 6,115,350,940 | 34,800 |
| 13 | non-specific-digest | 0 | 1 | 1M+15.9949 | 1,590,016,966 | 5,940 |
| 14 | non-specific-digest | 0 | 3 | 3M+15.9949, 3STY+79.966331 | 67,339,185,808 | 416,000 |
| 15 | non-specific-digest | 0 | 2 | 2M+15.9949, 2STY+79.966331 | 995,109,593 | 4,270 |

### 2.3 Database search

All database searches were conducted using the tide-search command. We used XCorr scoring with Tailor calibration (`--tailor-calibration T`) [10] using 8 CPU threads (`--num-threads 8`). The `precursor-window-size` and `mz-bin-with` parameters were set as specified in Table 1.

### 2.4 False discovery rate control

False discovery rate (FDR) estimates were calculated using the `assign-confidence` command in Crux. This tool implements a variant of the target-decoy competition (TDC) procedure [11], using decoy sequences generated automatically by tide-index. In this work, we use reversed peptides, with N-terminal and C-terminal amino acids left in place. Occasionally, the target contains two peptides that are reversed versions of one another; in such a setting, the corresponding decoys are created by shuffling rather than reversal. We have recently demonstrated that PSM-level FDR control is problematic due to dependencies between observed spectra [12]; therefore, in this work we focus on controlling FDR at the peptide level. The TDC procedure proceeds in three steps. First, the target and decoy PSMs produced by the tide-index command are sorted by score, from best match to worst match. Second, we eliminate from this sorted list any PSM whose corresponding peptide appears in a higher-ranked PSM, thereby producing a list of unique peptides. Third, for each target peptide in the list with score *τ*, we estimate the corresponding FDR as min (1, (*D*_*τ*_ + 1)*/T*_*τ*_), where *D*_*τ*_ is the number of decoy PSMs with scores greater than or equal to *τ*, and *T*_*τ*_ is similar, but for targets. The +1 correction is necessary to ensure valid FDR control [13], and the min() operator ensures that the FDR estimate does not exceed 1.

## 3 Results

### 3.1 Tide-index is fast and memory e?icient

In our first experiment, we demonstrate the improvement of the new tide-index in terms of execution time and memory consumption compared to the previous version. The analysis was carried out on a server equipped with an Intel Xeon CPU E5-2640 v4 2.40GHz processor with 20 cores, 128 Gb DDR RAM, and storage capacity of 10 TB operated by Ubuntu v18.04 OS. We selected a variety of tide-index parameter settings (numbers 1–5 and 11–14 from Table 3), which provided a series of indices of increasing size. Memory consumption was monitored with Ubuntu’s `time` command, setting the verbose parameter as well as the output file parameter, i.e. `/usr/bin/time -v -o time.txt`. The data recorded from the output file was the “Maximum resident set size.” We note that the previous version of tide index did not run to completion with parameter settings 12–14 because 128 Gb RAM was insufficient.

The results suggest that the new version of tide-index is approximately four times faster than the previous version (Figure 2A) and uses 2–3 times less memory (Figure 2B). The improved memory usage can be attributed to the reorganized data structures. We also tested the new `memory-limit` option by setting it to 32 GB. We note that most of the memory is used to ensure that only unique decoy peptides are generated and to ensure that the target and decoy peptides are distinct. Tide-index offers an option, called `--allow-dups T`, which allows the creation of duplicate peptides during decoy generation. Enabling this option additionally reduces the resource consumption by a large amount.

**Figure 2:**
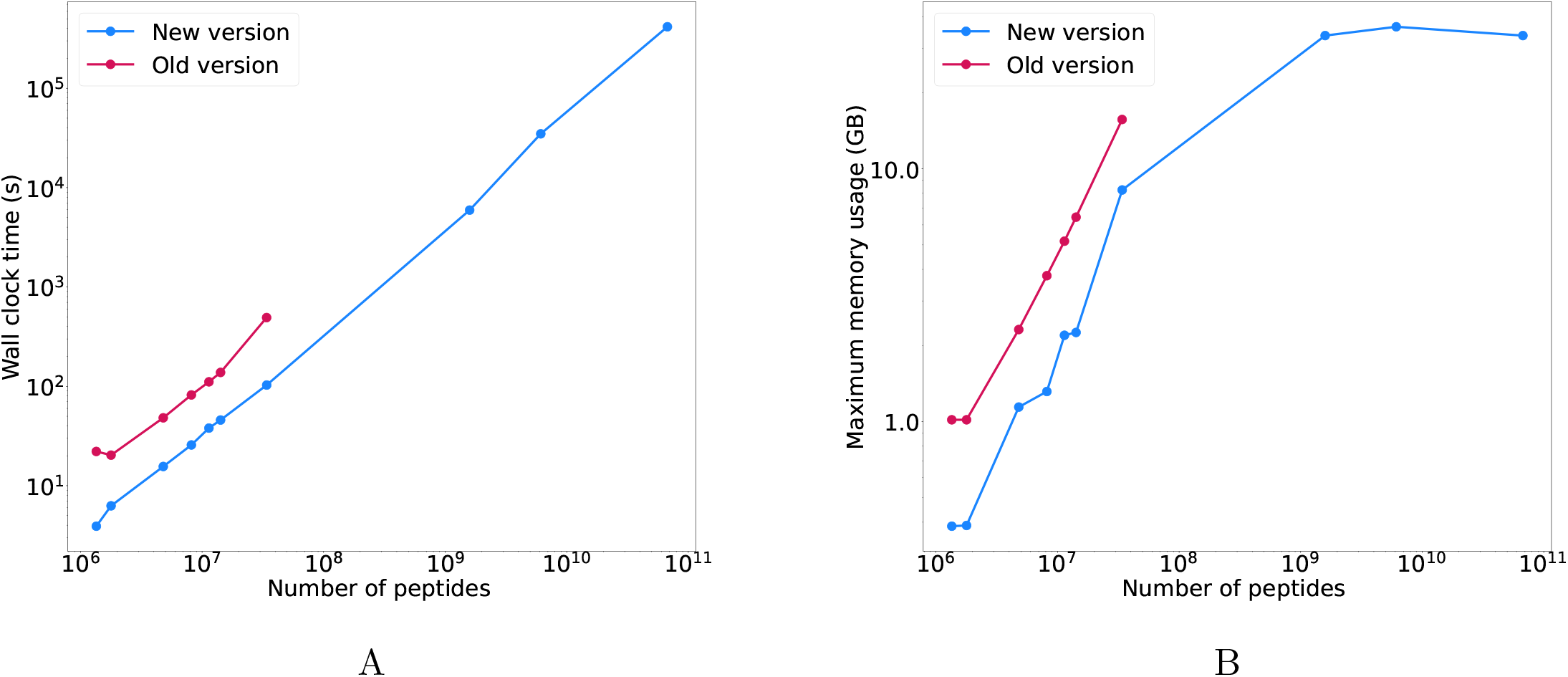
Speed and memory usage of tide-index. The figure plots (A) the wall clock time and (B) the maximum memory consumption required to produce an index of the human proteome as a function of the number of peptides in the index. The two series correspond to the old and new versions of tide-index. Points along each series correspond to the parameter settings 1-5 and 11-14 from Table 3.

The new version of tide-index allows us to create indices that would have been impossible to generate previously. For example, in our experiment, the largest peptide index was produced by the parameter setting #14, which created 67.3 billion unique peptides in nearly 5 days and which required 4.1 TB of disk space. Assuming that the resource consumption of the old tide-index is linear with respect to the number of peptides generated, as is suggested by Figure 2, then tide-index would have required around 10 TB of RAM to run to completion in around 20 days.

### 3.2 Parameter impact on database size

In our second analysis, we investigate how the parameter settings affect the number of the peptides generated. We generated a series of peptide index databases by varying three parameters.

First, we varied the digestion parameter and generated, for each proteome in our benchmark, three indexed databases using full tryptic digestion, partial digestion, and non-enzymatic digestion (parameter settings 10, 12, and 14 from Table 3), respectively. Changing this parameter increases the size of the peptide index by approximately a factor of ten, both when changing from full to partial digestion and when changing from partial to non-enzymatic digestion (Figure 3A). The fully digested indices contain between 82,719,463 peptides (yeast) and 300,423,422 peptides (human), whereas the non-enzymatic databases contain between 19,202,033,620 peptides (yeast) and 67,339,185,808 peptides (human).

**Figure 3:**
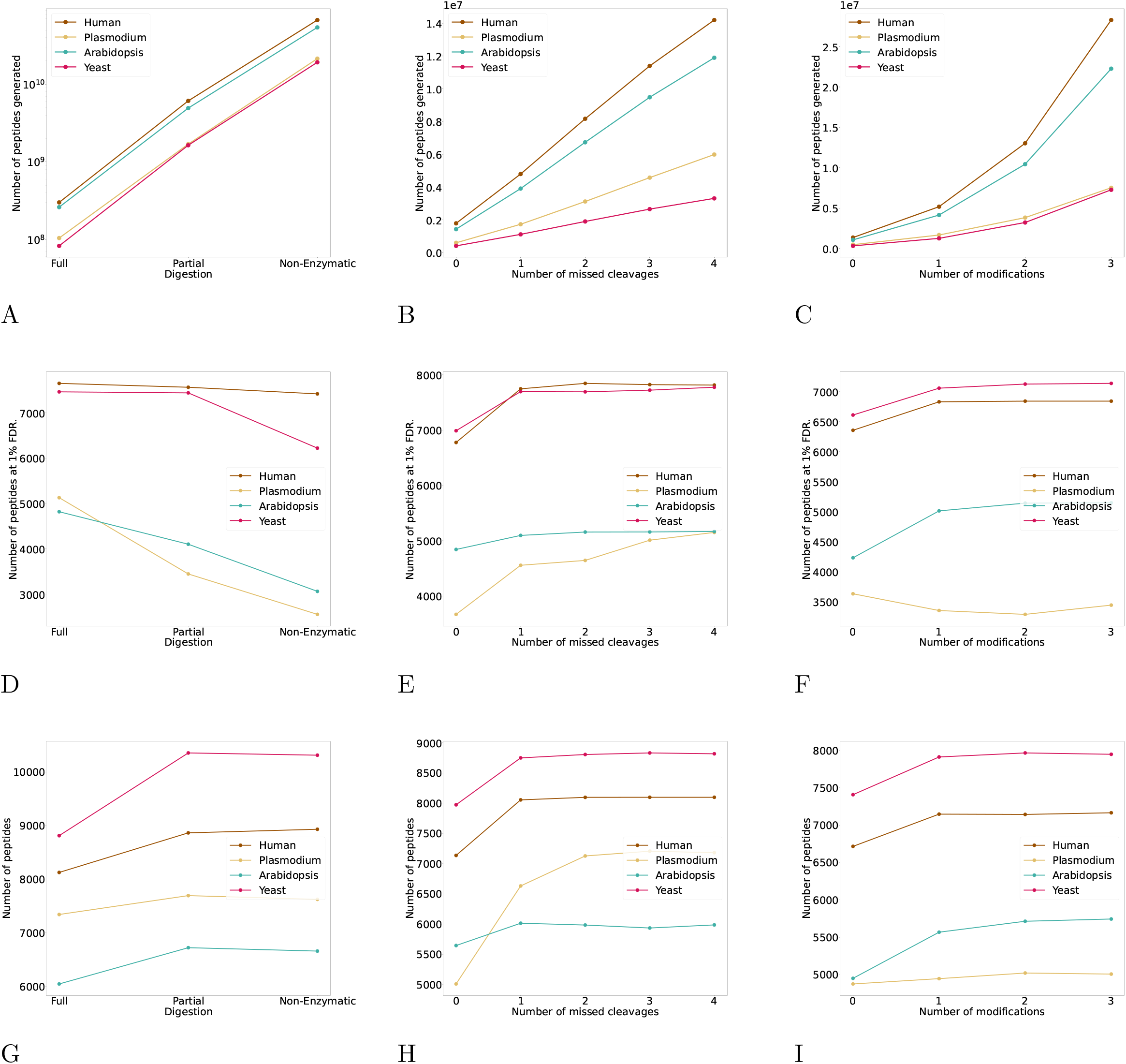
Parameter effect on database size and statistical power. (A–C) The figure plots database size as a function of (A) the type of digestion, (B) the maximum number of missed cleavages, and (C) the number of post-translational modifications. Each series corresponds to a different species. Panel (A) uses settings 10, 12, and 14 from Table 3; Panel (B) uses settings 2–6; panel (C) uses settings 1, 7, 8, and 9. (D–F) Similar to panels (A–C), except the y-axis plots the number of the peptides accepted at 1% FDR threshold, as computed using the Crux assign-confidence command. (G–I) Similar to panels (D–F), except that the FDR estimates are produced by Percolator rather than by assign-confidence.

Second, we varied the missed cleavages parameter from 0 to 4 (parameter settings 2–6) and observed a linear growth in the number of unique peptides (Figure 3B), though the gradient of the increase depends on the initial size of the database. On average, accepting an additional missed cleavage increases the size of the database by a factor of 2.9 for human but only 2.7 for yeast.

Third, we varied the number of variable modifications in the database. This was done by considering two different types of modifications—oxidation of methonine and phosphorylation of serine, threonine, or tyrosine—and then varying the total number of modifications allowed per peptide (from zero to three, corresponding to parameter settings 1, 7, 8, and 9). As expected, increasing the number of modifications results in exponential growth (Figure 3C).

### 3.3 Investigating the trade-off between database size and statistical power

Perhaps the most interesting analysis is to investigate the database size effect on the number of discoveries. In choosing parameters that affect the size of the peptide database, one faces a trade-off. On the one hand, limiting oneself to a small database increases the proportion of “foreign” spectra, that is, spectra whose generating peptides are not in the database and hence can never be identified. Essentially, the database is like a streetlight, and you will never find peptides that fall outside of the light. On the other hand, as the database gets larger the time to run the search increases proportionally and, more importantly, the statistical power to detect peptides decreases. The loss of statistical power arises because the high scores assigned to generating peptides begin to be overwhelmed by PSMs that score highly by chance, simply because such a large database was searched.

We thus investigated how the number of accepted peptides changes as we vary each of the three parameters: type of digestion, number of missed cleavages, and number of modifications (Figure 3D–F). Each search was carried out using Tide with Tailor calibration enabled, and decoy-based peptide-level FDR estimates were computed using the assign-confidence command in Crux. We observe that, for all four species, full tryptic digestion performs better than semi-tryptic, which in turn outperforms non-enzymatic digestion. On the other hand, for the other two parameters, allowing at least one missed cleavage or at least one modification seems to be beneficial in most cases (with the exception of *Plasmodium*, where even a single modification led to decreased performance). Furthermore, because these two parameters have a less pronounced effect on the size of the database, setting them to larger values (e.g., 3 or 4 missed cleavages) did not have a significant detrimental effect. And in one case (*Plasmodium*), 4 missed cleavages gave the best performance.

One caveat to the above analysis is that statistical power can depend strongly upon how the search results are post-processed. Accordingly, we further processed the Tide search results using Percolator, which uses a semi-supervised machine learning strategy to re-rank PSMs, and then uses decoy-based FDR estimation. The results (Figure 3G–I) suggest that using Percolator facilitates searching of larger databases. Most notably, in contrast to the results without a post-processor, in every case, switching from tryptic to semitryptic digestion yielded an increase in the number of detected peptides, with the most marked increase in the yeast dataset (an increase of 17%, from 8,812 to 10,312 peptides). Results for the other two parameters (Figure 3H–I) are qualitatively consistent with the previous results (Figure 3E–F), except that Percolator also removes the detrminental impact, for the *Plasmodium* dataset, of allowing too many modifications.

### 3.4 Tide can handle immunopeptidomics experiments

Finally, we demonstrate that Tide now is able to perform database searching for immunopeptidomics experiments. Previously, this type of analysis was impossible because this task involves the generation of non-specifically digested peptide databases which the old version of tide-index could not handle. In order to demonstrate that the new Tide can handle such tasks, we downloaded a randomly selected file from a recent analysis of peptides bound to class I major histocompatibility complexes (MHC) [9] (PXD017407 from Table 1). The peptide database was generated with non-specific digestion of the human proteome including oxidation of methonine and phosphorylation of S/T/Y as variable modifications and TMT-6plex labeling, and peptides were limited to 8–15 amino acid in length (see parameter settings 15 from Table 3). The resulting index contained nearly one billion peptides and required *∼*1.5 hours to generate. The database searching was performed with tide-search using standard parameters and with Tailor calibration [10], and the search results were evaluated with Percolator. This analysis detected 2,673 peptides at a 1% FDR threshold (Figure 4). These peptides overlap substantially (1624, or 93%) with the list of 1,753 peptides identified by Stopfer *et al*..

**Figure 4:**
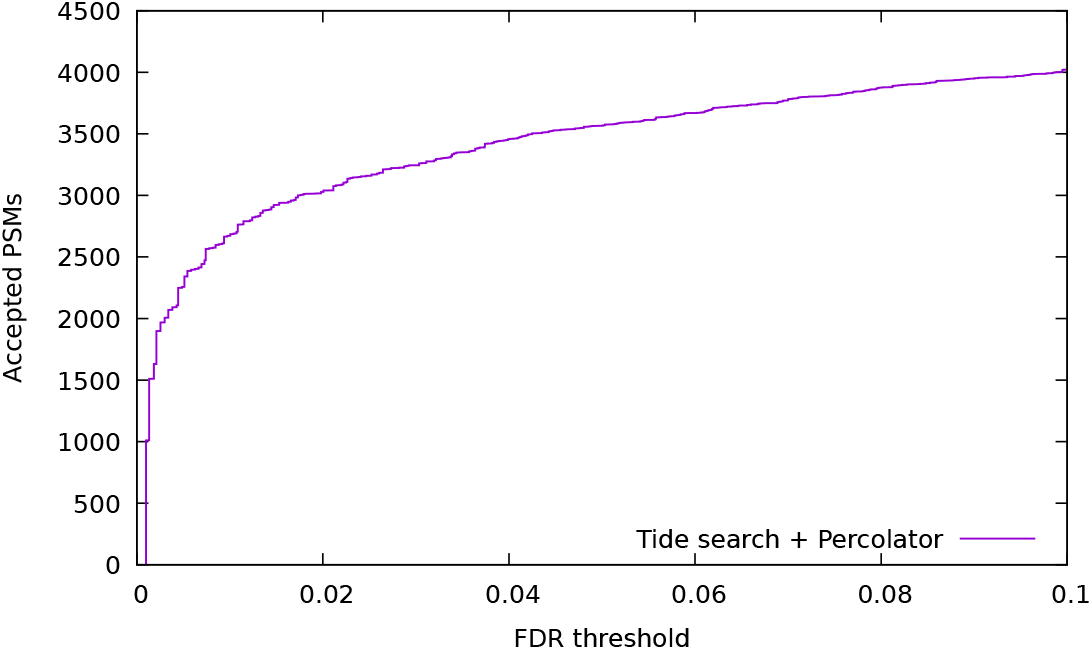
Crux analysis of MHC data. The figure plots the number of peptides accepted as a function of FDR threshold. The spectra are from a randomly selected file from PXD017407 [9].

## 4 Discussion

We have introduced an updated version of the Tide peptide database indexing step, which is able to efficiently handle very large peptide datasets, for application to metaproteomics or studies that aim to consider large numbers of post-translational modifications. We demonstrated that the new tide-index requires around three to five times less CPU time and memory than the previous version, and it can now generate as large peptide databases as the free disk space.

Our systematic analysis of the relationship between database size (i.e., total number of peptides) and several key parameters suggests several take-home messages. Notably, the type of digestion—full digestion, partial digestion, and non-enzymatic digestion—has the most profound impact on the size of the database, leading to approximately a 10-fold increase in database size for each transition. The effect of the other two parameters—number of missed cleavages and number of post-translational modifications—is smaller, though we did not extensively explore the latter parameter space. In particular, it seems clear that as the number of different types of modifications increases, the database size will likely increase dramatically. On the other hand, we also found that increases in database size were not always coupled with a loss in statistical power. In particular, we observe that including a few modifications (notably, oxidation of methionine, which is commonly induced during the mass spectrometry experiment) and allowing a few missed cleavages seems to generally be beneficial. Furthermore, setting these parameter values to be slightly too large does not appear to be problematic, as the number of detected peptides is fairly stable for larger values. Finally, it is notable that inclusion of a machine learning post-processor such as Percolator is helpful in combating the loss of statistical power induced by a large database. In particular, using partial digestion can be problematic for some studies if FDR control is carried out directly after the database search, but this is not the case in conjunction with Percolator. Many of the above observations are likely familiar to experts who regularly perform database search procedures, but we hope that the results presented here will be useful to those who want to better understand the potential impact of these parameters on their results.

On the other hand, as an empirical study of necessarily limited scope, this analysis leaves many questions unaddressed. In addition to more thoroughly exploring the space of possible post-translational modification parameters, one could imagine expanding this analysis to include a broader diversity of experimental studies, including more diverse proteomes. We hope that, if others aim to pursue such investigations, the updated version of Tide will provide a useful and efficient way to do that.

## Author contributions

FLNAA and AKF implemented the new tide-index. FLNAA carried out the experiments. WSN supervised the project. AKF and WSN wrote the manuscript. All authors read and approved the final manuscript.

## Availability

The new tide-index is available in the latest version of the Crux toolkit at https://crux.ms. Apache licensed source code is available, and pre-compiled binaries are provided for Windows, MacOS and Linux.

## Table of contents figure

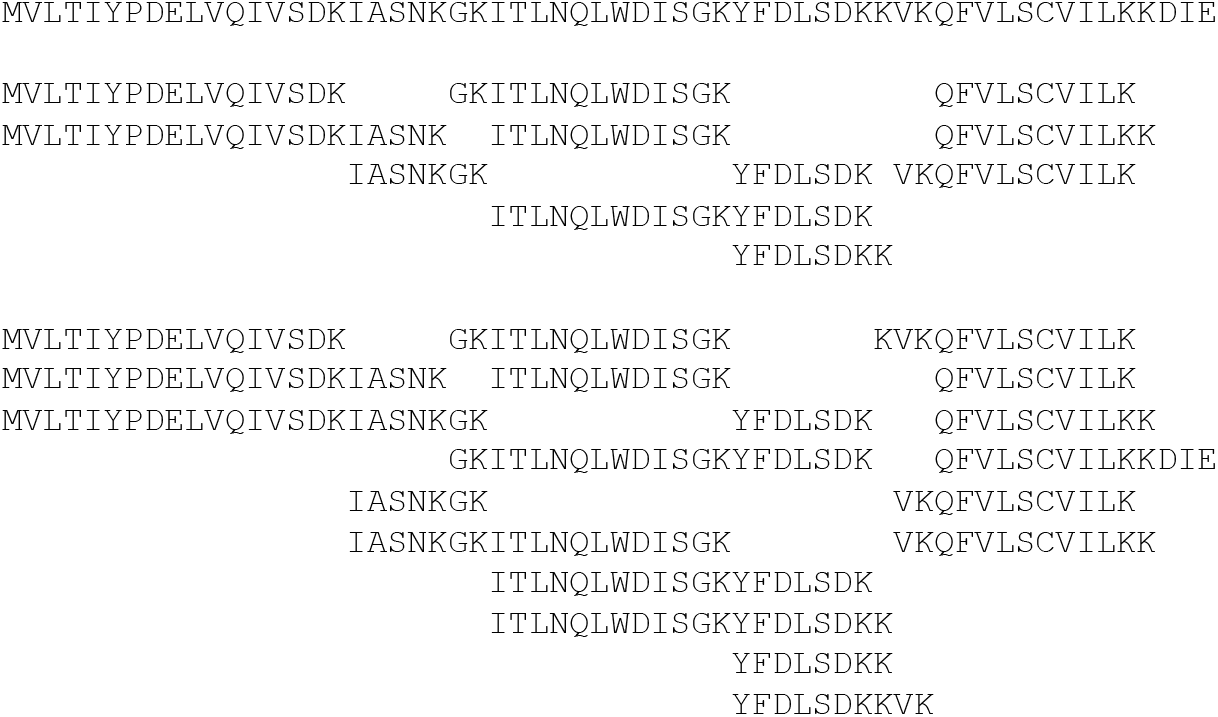

